# Optogenetic Locus Coeruleus Stimulation Improves Pupil Size Tracking of Cortical State

**DOI:** 10.1101/2024.02.02.578675

**Authors:** Evan Weiss, Yuxiang (Andy) Liu, Qi Wang

## Abstract

Brain state heavily influences our perception, cognition, and behavior. Multiple neuromodulatory systems, including the locus coeruleus – norepinephrine (LC-NE) system, contribute to the regulation of brain state. This research harnesses machine learning and optogenetics technologies to probe the impact of the LC on cortical state and pupillary responses in awake mice. Our integrative approach combines EEG recordings with pupillometry to capture the LC’s optogenetic activation, evoking notable EEG spectral power shifts synchronized with pupil dilation. These changes provide a noninvasive glimpse into the cortical states, modulated by the LC’s activity. Central to our methodology is the application of Support Vector Regression (SVR) modeling, which robustly correlates LC-induced pupillary changes with EEG power fluctuations. These insights help reinforce the LC’s crucial role in modulating prefrontal activity and its regulatory influence over arousal states. Beyond advancing our comprehension of the LC’s function, our work also highlights the potential for developing closed-loop stimulation systems. These systems, integrated with machine-learning techniques and multi-modal data, could offer precise therapeutic interventions for neurological disorders characterized by abnormal arousal states.

## I. Introduction

The locus coeruleus (LC), a noradrenergic brainstem nucleus, plays a crucial role in regulating many brain functions through the release of norepinephrine (NE) throughout the brain [1]. The LC’s heavy axonal projections to the prefrontal cortex are instrumental in regulating executive functions, including attention and behavioral flexibility [2]. While extensive research has explored the LC’s involvement in NE release across various cognitive paradigms, emerging studies suggest its heterogeneity might drive unique neural oscillatory dynamics, contingent on the prevailing cortical state [3].

The recording of cortical population neural activity, particularly in the frontal lobe regions, offers critical insights into global neural oscillations [4]. This method allows for a detailed examination of how spectral patterns are influenced and altered in response to various behavioral or neurotransmitter inputs, highlighting the dynamic nature of brain activity across different behavioral states [5]. By revealing these oscillatory changes across different cortical states, we gain a deeper understanding of the mechanism underpinning state transitions, often triggered by synchronized modulation within various neuromodulatory systems [6, 7]. Optogenetics offers a powerful tool for establishing real-time causality, enabling precise coordinated excitation of LC neurons, as well as unprecedented control of firing rate, through phasic or tonic stimulation patterns [8].

In this study, we investigated the influence of the LC on cortical neural oscillations by simultaneously recording cortical EEG and pupil size in awake, head-fixed mice, while optogenetically activating the LC. We achieved this by surgically implanting electrodes above the prefrontal cortex and delivering blue-light to Channelrhodopsin-2 (ChR2)-expressing LC neurons through an implanted optical fiber. Upon LC stimulation, we observed immediate and rapid pupil dilation, indicative of increased NE release and heightened arousal state. This physiological response was paralleled by a distinct shift in EEG spectral power, transitioning from low-frequency bands (Delta (1-3 Hz), Theta (4-7 Hz), Alpha (8-12 Hz), Beta (13-30 Hz)) to high-frequency bands (Gamma (30-100 Hz)) [9]. These findings align with previous arousal studies, where LC activation modulates these frequency bands, mirroring natural shifts in arousal and cortical state [10].

Next, we sought to use a machine learning approach to quantify the relationship between pupil dynamics and their corresponding EEG spectral changes, as both have been seen to serve as indirect measures of physiological arousal [11]. To achieve this, we employed a Support Vector Regression (SVR) model, a robust machine learning technique known for its effectiveness in handling high-dimensional data and nonlinear relationships. The SVR model was specifically utilized to build a decoder that characterizes the differences in EEG power changes during LC-evoked pupil dilations. This approach showcased the correlation between LC-evoked pupil dilations and EEG spectral changes, with an improving decoding performance as pupil dilation occurs. While providing evidence of the significant role the LC plays in modulating the pupil dilation response, it also helped uncover how the LC directly modulates specific EEG bands, seeming to be fundamentally different from spectral changes seen in naturally occurring, spontaneous dilations. Taken together, these results support the LC’s direct involvement in pupil-linked arousal, reinforcing its unique role in modulating cortical states.

## II. Methods

### A. Animal

All experimental procedures were approved by the Columbia University Institutional Animal Care and Use Committee (IACUC) and were conducted with compliance with NIH guidelines. Adult mice of both sexes (2 males and 1 females), aged 3 ∼ 7 months, were used in the experiments. The strains used were: Th-Cre (RRID: IMSR_JAX:008601) and DBh-Cre (RRID: ISMR_JAX:033951). All mice were kept under a 12-hour light-dark cycle.

### B. Surgical procedures

Animals were anesthetized under isoflurane (5% induction, 2% maintenance) and fixed in a stereotaxic frame. Body temperature was maintained at 37 °C, using a feedback-controlled heating pad (FHC, Bowdoinham, ME). Once the animal’s condition stabilized and before an incision was made, lidocaine hydrochloride and buprenorphine (0.05 mg/kg) were administered subcutaneously. After surgery, Baytril (5 mg/kg) and Ketoprofen (5mg/kg) were administered, as well as subsequently four additional times after each surgery day. Animal’s weight was measured once a day for 5 days. For adeno-associated viral vector (AAV) injections, burr holes were drilled above the LC, with saline applied to each craniotomy to prevent drying out of brain surface. Pulled capillary glass micropipettes (Drummond Scientific, Broomall, PA) were back-filled with AAV solution and injected into the target brain regions at 0.7nL/s using a precision injection system (Nanoliter 2020, World Precision Instruments, Sarasota, FL). The pipette was left in place for at least 10 minutes between injections and slowly withdrawn. To optogenetically activate the LC, pAAV-EF1a-double-floxed-hChR2-(H134R)-mCherry-WPRE-HGHpA (ChR2) was injected into the LC (AP: -5.3 mm, ML: 0.85 mm, DV: -3 mm). Immunohistology staining for tyrosine hydroxylase (TH) antibodies revealed a prominent expression of mCherry-tagged ChR2 in TH-positive neurons in the LC (**Figure 1C**). After injection, an optical fiber (200μm diameter & NA = 0.39) was implanted with the tip of the fiber placed approximately 0.15 mm above injection site. C&B Metabond (Parkell Inc., Edgewood, NY) was used to build a headcap to bond the fibers and the head bar. Three EEG recording screws (NeuroTek-IT Inc, Toronto, Canada) were gently threaded into the skull to touch the prefrontal cortex and occipital lobe (grounding). A custom-made EEG head stage was prepared prior to each implantation, consisting of a solderable bread board to thread each EEG screw wire. Ferrules and headplate were cemented in place with dental acrylic (Prime Dental Manufacturing, Chicago, IL). All recording was performed 3-4 weeks following surgery to allow enough time for viral expression (**Figure 1B**).

**Figure 1.**
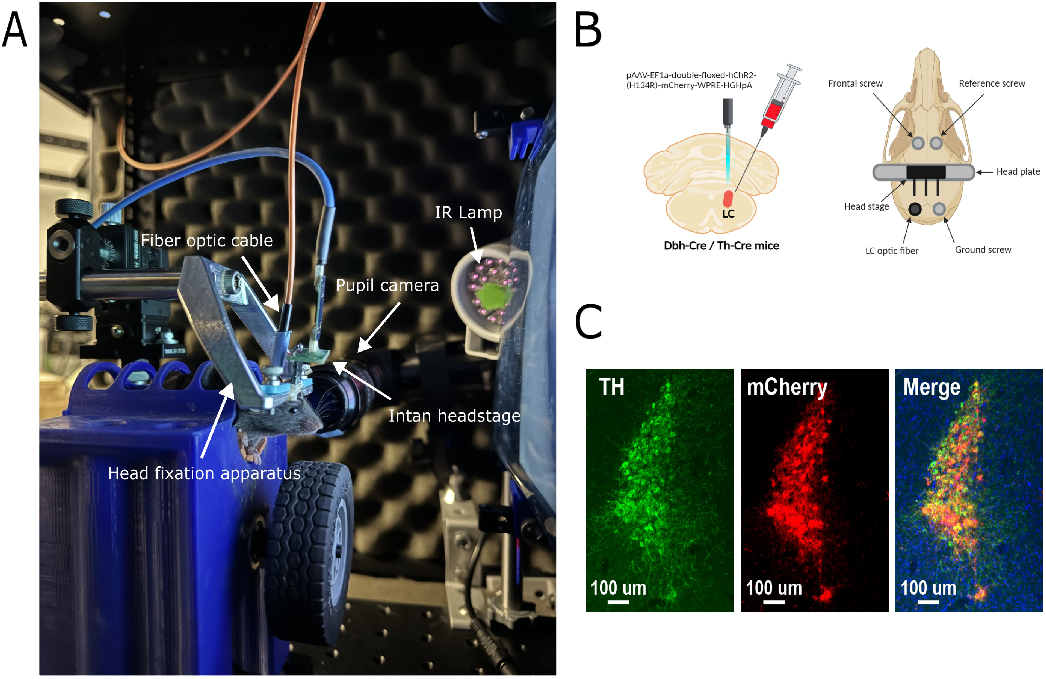
Surgical Methods and Recording Setup. A) Picture inside custom designed recording rig, equipment enabling real-time synchronized acquisition. B) Visualization of viral injection and optical fiber in the LC (left), top-down view of skull, bone screws (EEG recording) & headplate/ headstage for recording (right) created with BioRender.com. C) Histological verification of TH+ neurons (green) coexpressed with mCherry ChR2 (red) with DAPI (blue).

### C. Real-time Stimulation and Recording System

We used a blue laser (λ = 473 nm, PlexBright LED Module, Plexon Inc., Dallas, TX) and a fiberoptic cable for optogenetic stimulation of the LC. A RHD USB interface board (Intan Technologies) was connected to a 16-channel recording head stage (Intan Technologies) and soldered to a custom-made PCB breakout board, for easy plug-in data acquisition to the head stage on each mouse. For real-time data acquisition and TTL signal alignment, a Bpod State Machine (Sanworks) was used with custom Python experimental scripts to synchronize optogenetic stimulation events, EEG recording, and pupil camera acquisition. For pupillometry recordings, a USB camera (BFS-U3-16S2M, Point Grey) was triggered by a 10 Hz TTL from the Bpod. In each recording, the mouse sat in a 3D printed head-fixation platform (**Figure 1A**). Every 30 seconds, an optogenetic stimulation with a randomly chosen stimulation frequency (0 Hz, 10 Hz, or 20 Hz; 10ms-ON, 90ms-OFF) was delivered for a 2-second duration. The DeepLabCut (DLC) toolbox was used to segment the pupil contour in a pre-defined region of interests (ROI). The resnet50 deep network was trained on each labeled frame and analyzed each pupil video. Circular regression was applied, and subsequently uploaded to MATLAB for further analysis. All subsequent neural and stimulation data was extracted and analyzed in Python and MATLAB. EEG data was sampled at 1 kHz and collected with a 60 Hz notch filter.

## Results

### A. LC stimulation evokes pupil dilation

We imaged the pupil of 3 awake, head-fixed mice in response to optogenetic LC stimulation. Upon administering LC stimulation, we consistently observed an immediate and marked pupil dilation (**Figure 2A**). We used DLC to quantify pupil size and found a significant difference (p<0.005) between pre-stimulation and 2 seconds post-stimulation onset across all subjects in stimulation trials. Conversely, in non-stimulation trials, there were no significant changes in pupil size post-onset (p= 0.85) (**Figure 2B**).

**Figure 2.**
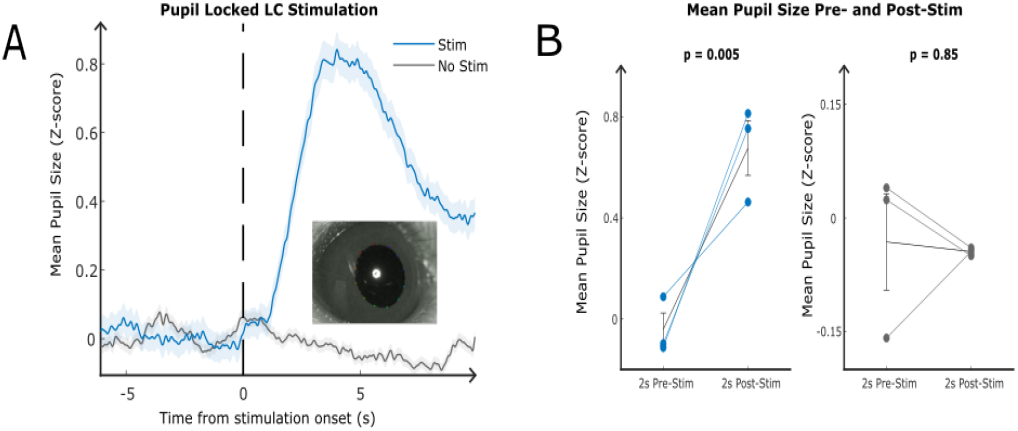
Pupil LC stimulation response A) Mean pupiI size (z-scored) locked to LC stimulation onset vs non-stimulation trials & visualization of labeled pupil frame with DeepLabCut. B) Mean pupil size change to stimulation pre-stim and post-stim.

### B. Cortical state shifts triggered by LC stimulation

During LC stimulation, we conducted simultaneous EEG recordings to our pupil size measurements. Prior to band-specific analysis, we calculated the power spectral density for all mice during both the 2-second stimulation and non-stimulation periods, revealing a significant shift from low-to high-frequency power (**Figure 3A**). Visualizing our spectral changes using a stimulation-locked spectrogram showcases a similar trend (**Figure 3B**). Further examination of these power changes across each frequency band revealed distinct patterns: the lower bands (Delta (1-3 Hz), Theta (4-7 Hz), Alpha (8-12 Hz) displayed a transient decrease in power during the rise of pupil dilation, followed by a rapid return to baseline after the stimulation period (**Figure 3C**). Conversely, higher frequency power (65-100 Hz), showed an inverse relationship, increasing to a local maximum at the peak of pupil dilation, then gradually declining to baseline levels.

**Figure 3.**
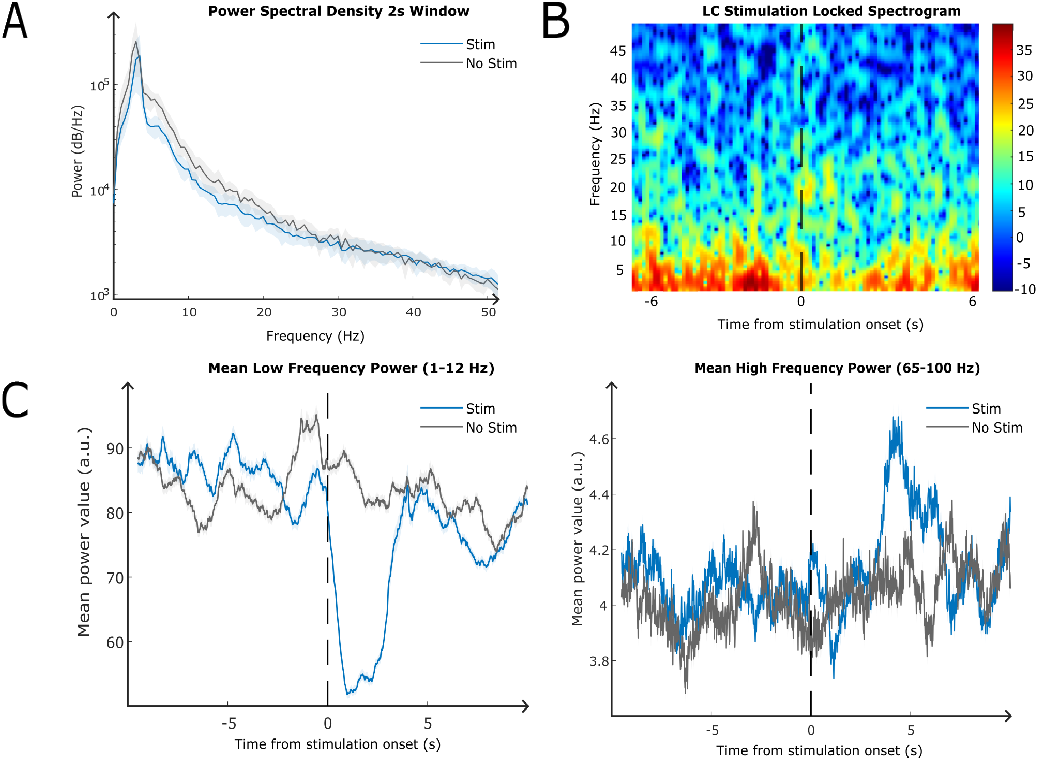
EES LC stimulation response. A) Power Spectral Density plot of stimulation vs no stimulation period (2 seconds). B) Spectrogram locked to LC stimulation response C) Mean PSD power locked to stmulation onset, low frequency power (left) high frequency power (right)

### C. SVR Model for predicting EEG power from pupil size

In order to further classify how the activation of the LC can orchestrate different EEG power changes, we looked to machine learning, specifically Support Vector Regression (SVR) to understand how accurately pupil size can track changes in EEG band power. Given the difference in sampling rates of our pupil (10 Hz) and EEG (1 kHz) data, we first up-sampled our pupil size data to 100 Hz, using spline interpolation. To track the fluctuation of power of different frequency bands, we divided the average power within each 20-second period into 2000-time windows. This was achieved by computing the area under the curve across each frequency band in the power spectral density (PSD) function. Applying a 1-second moving window with a 0.01-second shift across the EEG trace, now resulted in each window having a 2000-point vector. Each point here represented the average PSD power for that specific 1-second moving window (**Figure 4A**).

**Figure 4.**
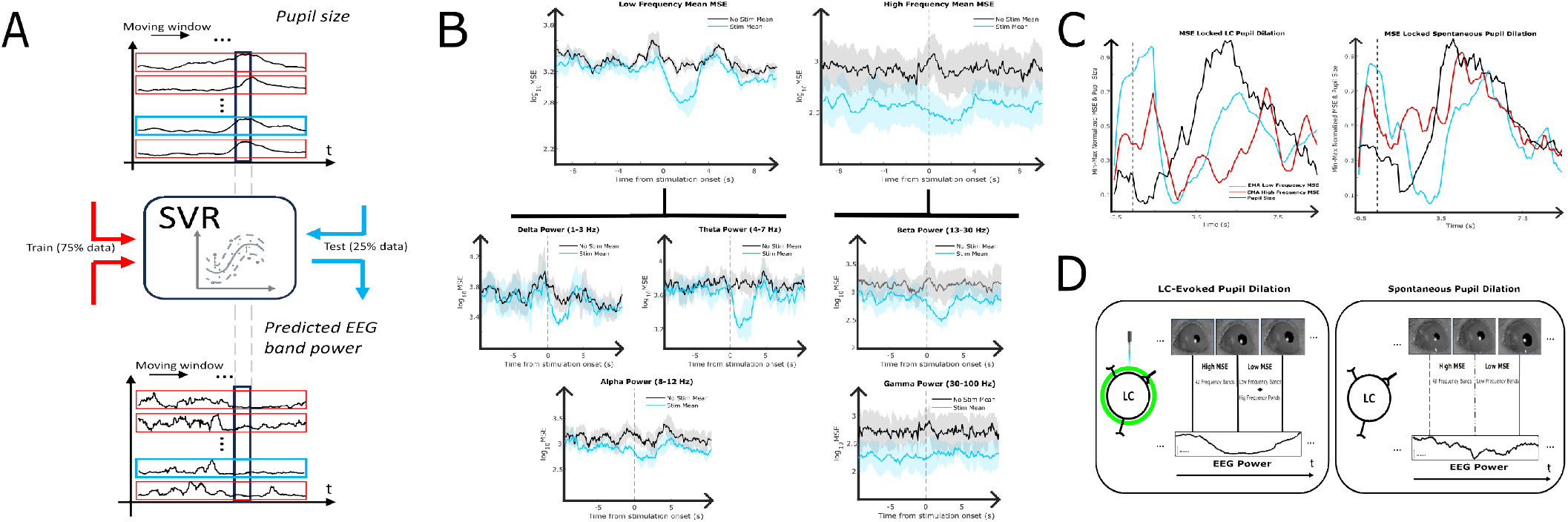
Support Vector Regression Modeling. A) Visualization of SVR model, train/test split and moving window segmentation of EEG and pupil traces. B) Mean MSE across time between stimulation and no stimulation traces across low & high frequency bands (top), and each individual band (Bottom).C)MSE locked to pupil dilation Between LC evoked(left)and spontaneous pupil dilations (right), visual difference in low vs high frequency MSE across time. D) Visualization of MSE changes across EEG frequency bands differing between LC-evoked and spontaneous pupiI dilations, orchestrated through the degree of LC activation.

The newly split train and test data was passed into a sigmoidal kernel SVR model individually trained to each subject. Mean squared error (MSE) was extracted from each 0.01-s moving window, allowing a glimpse into how prediction accuracy fluctuates over the entire 20-s period. Next, we visualized the mean MSE for each frequency band, aligning low- and high-frequency together, as well as each sub-band to our stimulation onset (**Figure 4B**). Here we saw notable drops in mean MSE for Delta, Theta, Alpha, and Beta power, but an increase for Gamma, approaching minimums during the onset of pupil dilation. Mean MSE drop from baseline showcases a drop of 10% in MSE for Delta, Theta, and Beta power, a 5% drop for Alpha power, and a 5% increase in Gamma power. This pattern aligned with the observed decrease in EEG power and increase in pupil size during LC-induced dilation. Aligning our mean MSE for low- and high-frequency bands back to our model’s pupil dilation segments, allowed for higher resolution into how the SVR’s predictive capability changes across our input data. Here we looked at the difference between an LC-evoked dilation and a spontaneous dilation’s (extracted as random non-stimulation pupil dilations) mean MSE. Surprisingly, only the high-frequency power from LC-evoked dilations converged with the mean low-frequency MSE, while spontaneous dilations lacked a similar high-frequency convergence (**Figure 4C**). Spontaneous dilations seem to lack the degree of beta wave modulation caused by direct LC activation, potentially implicating the involvement of other neuromodulatory systems and adding complexity to the LC’s effect on cortical state (**Figure 4D**). These findings highlight the efficacy of our SVR model in decoding pupil into EEG power, highlighting a unique role of the LC in modulating cortical states, and pupil dilations.

## IV. Discussion

In this study, we have integrated a robust engineering framework to concurrently capture cortical state alterations, pupil dynamics, and selective optogenetic stimulation in an awake, head-fixed mouse model. The primary objective of this study was to elucidate the role of the LC in modulating cortical states as evidenced by distinct spectral power shifts within EEG activity, and to quantitatively link pupil size changes to cortical states. Our findings demonstrate that LC optogenetic activation precipitates not only a rapid and sustained pupil dilation, but also induces a pronounced transition in EEG spectral power from lower to higher frequencies. This dual response pattern mirrors the observations documented in human research, wherein similar power shifts in Alpha and Theta bands are discernible in response to deep-brain or vagal nerve stimulation, accompanying changes in pupil size [12, 13]. The predictive capacity of our Support Vector Regression model further substantiates the correlation between pupil size during LC stimulation and the preferential modulation of EEG low-frequency power (Delta, Theta, Alpha, Beta) relative to high-frequency bands (Gamma). Given the direct nature of optogenetic stimulation, we infer a pivotal role of the LC in mediating these cortical state dynamics. This is indicative of the LC’s unique function in a spontaneous vs a stimulation-evoked dilation. With LC-evoked enabling a higher predictive ability in both low- and high-frequency bands aligning with pupil dilation. Spontaneous dilations lack the convergence of high frequency MSE, implicating the potential activity of other neuromodulatory systems or brain regions during a pupil dilation.

Looking ahead, the implications of our results extend into the domain of neurological therapeutics and diagnostics. The ability of the LC the modulate both autonomic and cortical response underscores its potential as a target for interventions in disorders characterized by dysregulation of arousal and attentional pathways. Furthermore, the use of SVR models for predicting EEG spectral power based on pupil size opens up new avenues for real-time stimulation and monitoring systems. Closed-loop stimulation systems could leverage the power of non-invasive data modalities such as real-time pupil or EEG to gain insights into the activity of deep brain structures like the LC, as well as performing targeted stimulation to optimize a specific cortical state.

## V. Conclusion

In conclusion, our investigation confirms the LC as a critical nexus in the regulation of arousal and cortical states, mediated through pupil dynamics. The LC’s ability to orchestrate EEG spectral power shifts has profound implications on the neural underpinnings of arousal and attention. By leveraging machine learning, we have enhanced our capacity to decode and predict these shifts, showcasing a unique role of the LC in shaping cortical state. Thus, our work not only delineates the LC’s pivotal role in regulating cortical oscillations but also sets the stage for future advances in closed-loop stimulation systems for neurological therapeutic interventions.

## Acknowledgments

This work was supported by the Air Force Office of Scientific Research under award number FA9550-22-1-0337. Any opinions, findings, and conclusions or recommendations expressed in this material are those of the authors and do not necessarily reflect the views of the United States Air Force.

## Notes

### Competing Interest Statement

The authors have declared no competing interest.

